# The effect of long-term management on wild pig (*Sus scrofa × domesticus*) populations across the southeastern United States

**DOI:** 10.64898/2026.04.16.719012

**Authors:** John R. Foster, Kim M. Pepin, Ryan S. Miller

**Affiliations:** United States Department of Agriculture, Animal and Plant Health Inspection Service, Veterinary Services, Center for Epidemiology and Animal Health, Fort Collins, CO, USA; Oak Ridge Institute for Science and Education, United States Department of Energy, Oak Ridge, TN, USA; United States Department of Agriculture, Animal and Plant Health Inspection Service, Wildlife Services, National Wildlife Research Center, Fort Collins, CO, USA

## Abstract

1. The management of invasive species often emphasizes removals to manage populations. However, evaluating the success of this management technique remains challenging, especially at large scales. Understanding the relationship between removal intensity and population growth is essential for determining when management achieves desired outcomes.
2. We used management removal data (removal resources [e.g. trapping] and relative effort [trap nights]) to estimate population density, demographic structure, and growth rates of invasive wild pigs (*Sus scrofa× domesticus*) across a large landscape. From the management data and population estimates, we inferred population trajectories in the absence of removals and quantified the proportion of the population removed by the most widely used methods to control wild pigs. We then compared observed removal intensities and population growth rates to predict expected population trajectories immediately after management occurs.
3. Results suggest substantial spatial and temporal variation in wild pig growth rates and variation in the effectiveness of removal efforts. Additionally, removing wild pigs at higher densities had a greater effect on limiting population growth than removals conducted at lower densities, though both are important. However, on large properties, removal intensity was often insufficient to offset population growth, indicating that management effort does not scale to large areas.
4. These results demonstrate how removal data and population modeling can provide robust inference on population dynamics and management effectiveness, offering a scalable framework for evaluating and improving invasive species control programs. We also discuss the current limitation of how effort is defined for different large-mammal removal techniques, and offer potential solutions for a more complete definition, such as going beyond trap nights and including constraints on personnel, equipment, and logistics.

## Introduction

Invasive species are among the most pervasive drivers of ecological change; they can reshape community structure, alter nutrient cycling, and threaten biodiversity across ecosystems worldwide (Crowl et al. 2008; Didham et al. 2005; Pyšek et al. 2020; Simberloff et al. 2013; Vitali et al. 2023). Management of invasive species often centers on lethally removing individuals with the intent to reduce or eliminate populations and mitigate ecological and economic impacts (Epanchin-Niell and Hastings 2010). Yet the success of such interventions is rarely evaluated in terms of their effects on population dynamics, especially at large geographic extents (Green and Grosholz 2021). For removal-based management to be effective, population control efforts must reduce abundance to levels that achieve desired management outcomes (Prior et al. 2018), otherwise known as functional eradication (Green and Grosholz 2021).

Understanding whether management achieves a desired outcome, such as population reduction, requires knowing the intensity of removal relative to population size, which is necessary for identifying thresholds of removal required to reduce abundance (Pepin et al. 2023; Hone 2001). Furthermore, thresholds at which population decline or elimination could occur must be estimated in relation to a population’s growth rate (Hone 2001; Hone et al. 2010; Manlik et al. 2022; Pepin et al. 2023). Additionally, compensatory mechanisms such as increased survival, reproduction, or immigration may maintain or even elevate population size in response to removal efforts (Hanson et al. 2009; Walsworth et al. 2020; Zipkin et al. 2009). Therefore, effective invasive species control requires understanding the link between population growth rates and density reduction, which is vital for preventing spread (Pepin, et al. 2017; Pepin et al. 2023).

Advances in demographic modeling have shown that it is feasible to estimate population density and demographic structure using removal data alone, particularly when removals occur systematically over space and time (Davis et al. 2016, 2022; Taylor et al. 2025; Foster et al. 2025). By incorporating assumptions about detection, respective vulnerability to various removal methods, and life history, removal data can be used to infer not only current population size but also likely population trajectories in the absence of management (Udell et al. 2022; Botterill-James et al. 2024). Demographic modeling enables managers to separate the effects of removal from natural population trends and assess whether observed declines in abundance result from management or demographic processes.

This inferential power is especially relevant for wild pigs (*Sus scrofa× domesticus*), a highly invasive species that causes extensive ecological and economic harm (McKee et al. 2025; Barrios-Garcia and Ballari 2012; VerCauteren et al. 2024). Their high fecundity, generalist behavior, and capacity for rapid recolonization make wild pigs particularly difficult to control (Garabedian and Kilgo 2024). Although wild pig management programs have occurred broadly over the last several decades, where individuals are removed primarily through trapping, aerial gunning (i.e. from an helicopter of fixed wing platform), and ground-shooting, few programs assess whether these efforts translate into sustained population declines (Snow et al. 2025). In other words, without knowing what proportion of the population is being removed or population growth rates in the absence of intervention, managers cannot determine whether their efforts are ecologically and/or economically effective (Davis et al. 2018).

Critically for wild pigs and other invasive species, these assessments must occur at large spatial and temporal scales to capture their dynamic and heterogeneous nature, which can be difficult because many wildlife management programs operate with limited budgets (White et al. 2022). Population growth rates, immigration, and removal efficiency all vary across landscapes and seasons, and short-term or localized studies may miss broader trends in regulation (Snow et al. 2024, 2020). Furthermore, wild pig populations may exhibit compensatory growth in response to removal pressure (Servanty et al. 2011; Miller et al. 2023), and their population growth rates can be highly reactive (Tabak et al. 2018). By quantifying the proportion of the population removed across multiple years and regions, and comparing these rates to model-based estimates of the removal needed to achieve a specific management goal, we can better understand the conditions under which removals lead to population reduction or elimination (Zipkin et al. 2009; Davis et al. 2022).

In this study, we use systematic removal data to estimate wild pig population density, demographic structure, and population trajectory across a large landscape. Our goal is not to find an optimal control strategy for wild pigs, but to link removal intensity to population growth rates, improve inference about management effectiveness, and inform theory-based strategies for invasive species control. Specifically, we aim to (1) quantify the proportion of the population removed through control efforts and evaluate how variation in this proportion affects population outcomes over space and time; (2) quantify the variation in wild pig population growth rates over a large geographic extent; (3) estimate the density increase prevented in the absence of management; (4) and determine the effort needed to achieve certain management goals such as regulation and suppression.

## Methods

The model described in Foster et al. (2025) was initially used to predict population density from wild pig removal data that was obtained from the United States Department of Agriculture – Animal Plant Health Inspection Service – Wildlife Services (USDA-APHIS-WS). USDA-APHIS-WS maintains a database that contains records of all management actions, including the management method used (e.g. traps, snares, ground-shooting, and shooting from fixed wing aircraft or helicopters), relative effort (e.g. flight hours, trap nights), and the number of wild pigs removed. Initially, the integrated removal model (hereafter “removal model”) in Foster et al. (2025) was used to estimate wild pig density at 105 properties where USDA-APHIS-WS conducts wild pig management. The removal model estimated abundance given removal effort, the method used for removal, county-level landscape features, survival, and per-capita recruitment. The ecological process was dynamic, meaning the previous density estimate (i.e. at time t-1) was used to estimate current density (i.e. at time t). The model used a 28-day primary period, binning all removal actions within the primary period as the units of removal effort and assuming population closure during that interval (Link et al. 2018).

We iteratively fit the Foster et al. (2025) model using Bayes’ theorem to get density estimates for the properties with suitable data (n=995) where USDA-APHIS-WS conducts management activities. Initial model runs from Foster et al. (2025) yielded a posterior probability distribution that then served as the prior distribution for subsequent model runs (Ellison 2004). Given the size and complexity of the dataset, we employed an iterative approach to density estimation as fitting the removal model to all 995 properties at once was infeasible. Specifically, removal effort was documented in 12,432 distinct primary periods. However, because the removal model was dynamic (predicting density for every primary period in each property, first to last, including unsampled ones), there was a total of 35,472 primary periods density was predicted. This means that only 35% of primary periods were observed, rendering a single model fit impractical due to the significant data sparsity in these data. See appendix S1.3 for further details on this approach.

### Population growth

Realized population growth rate (𝜆_𝑖𝑡_) was calculated for each primary period 𝑡 in each property 𝑖. Removing individuals was essential to the model meaning we incorporated the number of individuals removed (i.e. catch, 𝐶_𝑖𝑡_) when calculating growth rate as our model assumed that growth occurs after removals (Skalski et al. 2005). Specifically,

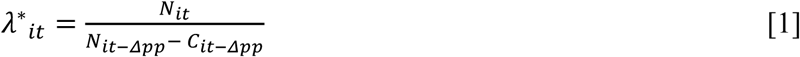

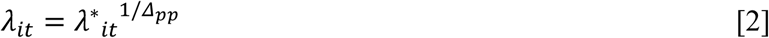

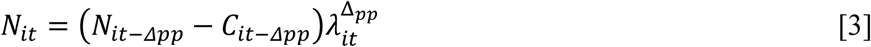

where 𝜆^∗^is the apparent growth rate between observed primary periods (eq. 1), and 𝛥𝑝𝑝 is the number of primary periods between observations. The instantaneous population growth rate per 28-day period (𝜆_𝑖𝑡_, eq. 2) can then be used to estimate abundance (𝑁) across missing time points (eq. 3). We then estimated the mean growth rate for each property (i.e. population) as 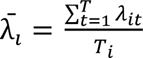 where 𝑇 is the number of observed primary periods in property 𝑖.

### Density prevented without management

To assess the effect of management in terms of density prevented (i.e. predict density if management did not occur), we ran simulations of the population model (Appendix 1 Section 1.1) where the number of individuals removed was set to zero and compared the simulations to posterior density estimates. We ran 10,000 simulations, where initial abundance was drawn randomly from the set of estimated abundances (e.g. posterior median abundances). We ran each simulation for five years, and within each simulation we ran 1,000 ensembles, where each ensemble corresponds to a set of demographic parameters that were chosen at random, with replacement, from the joint posterior of the final fit to data. Each simulation used the same set of randomly drawn parameters. A simulation was discarded if the initial timepoint was at the end of a properties’ time series (because there would be nothing to compare it to), or if the initial abundance was greater than carrying capacity, which was set to 30 individuals/km^2^ (Pepin, et al. 2017).

### Expected population trajectory

The concept of the Expected Population Trajectory (EPT) is crucial for understanding population dynamics and effects of management interventions. The fundamental distinction between EPT and the actual proportion of the population removed (APR) lies in how they account for the inherent growth rate of a population. APR is a metric that represents the direct reduction in population size due to a specific removal event. It is a straightforward measure of the immediate impact, without considering the population’s natural ability to grow or decline. For instance, if 50 individuals are removed from a population of 100, the APR is 0.5 (50/100). The expected population trajectory (EPT), by contrast, provides a more nuanced understanding by integrating the population’s intrinsic growth rate. It projects how the population is expected to change in relative terms after accounting for both removals and natural growth and is a more accurate assessment of the net effect of APR on the population’s immediate future size. EPT is calculated as

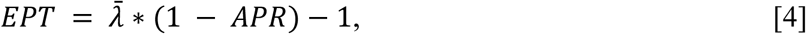

the derivation of which can be found in Appendix 1 Section 1.2. If 𝐸𝑃𝑇 = −1, the population is expected to be eradicated, as all individuals have been removed (i.e. 𝐴𝑃𝑅 = 1). If 𝐸𝑃𝑇 = 0, the population is expected to have no net change in size, and then equation 4 can be rearranged to 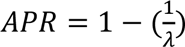, and represents the threshold APR needed to reduce a population (Hone 2001). This occurs either with no removals and a stable population, or when removals are precisely balanced by the population’s growth rate and signifies a dynamic equilibrium where gains (i.e. births/immigration) are offset by losses (i.e. deaths/removals/emigration). If 𝐸𝑃𝑇 = 1, the population is expected to double and occurs when the reduction effect of APR is significantly outweighed by the population growth rate.

## Results

### Population estimates

Here, and throughout the manuscript, we describe distributions by their median and central 90% interval (i.e. CI; the 5% and 95% quantiles). Across the 12 states and 11 years where USDA-APHIS-WS removals occurred (n=12,432 individual primary periods from 2014-01-28 to 2024-06-30 across 995 properties), the distribution of median posterior density estimates had a median of 1.28 (0-16.25) wild pigs/km^2^. There was a consistent downward trend in density through time. In 2014 the median density was 2.26 (0.08-17.35) wild pigs/km^2^, and by 2024 median density was 0.54 (0-14.25) wild pigs/km^2^ (Appendix 1 Figure S5). Note that 2024 was an incomplete year of sampling. Furthermore, these are not national scale density estimates or trends, but rather a summary of estimated densities at properties where USDA-APHIS-WS conducted removals over this time period.

### Proportion of the population removed varies by method

Removal methods were not used at all properties or even in all states. Additionally, resource constraints and environmental conditions can dictate which methods are used and when. Therefore, understanding the effectiveness of context-specific management is critical for evaluating future removal efforts.

There were nine states in which the median proportion of the population removed in a 4-week period by traps was zero (Figure 1), and the maximum came from Texas with a median of 0.05 (0-0.35). For snares, there were three states where the median proportion of the population removed was zero, and the maximum came from Louisiana with a median of 0.16 (0.15-0.17). For ground-shooting, the range in the median proportion of the population removed was between 0.03 (0.02-0.63) and 0.42 (0.36 - 0.49) for Oklahoma and South Carolina, respectively. There were two states that used fixed wing aircraft, which removed a proportion between 0 (0-0.02) and 0.01 (0-0.3) for Oklahoma and Texas, respectively. Lastly, for helicopters, the range in the median proportion of the population removed in a 4-week period was between 0.04 (0-0.07) and 0.42 (0.11-0.72) for New Mexico and Mississippi, respectively.

**Figure 1.**
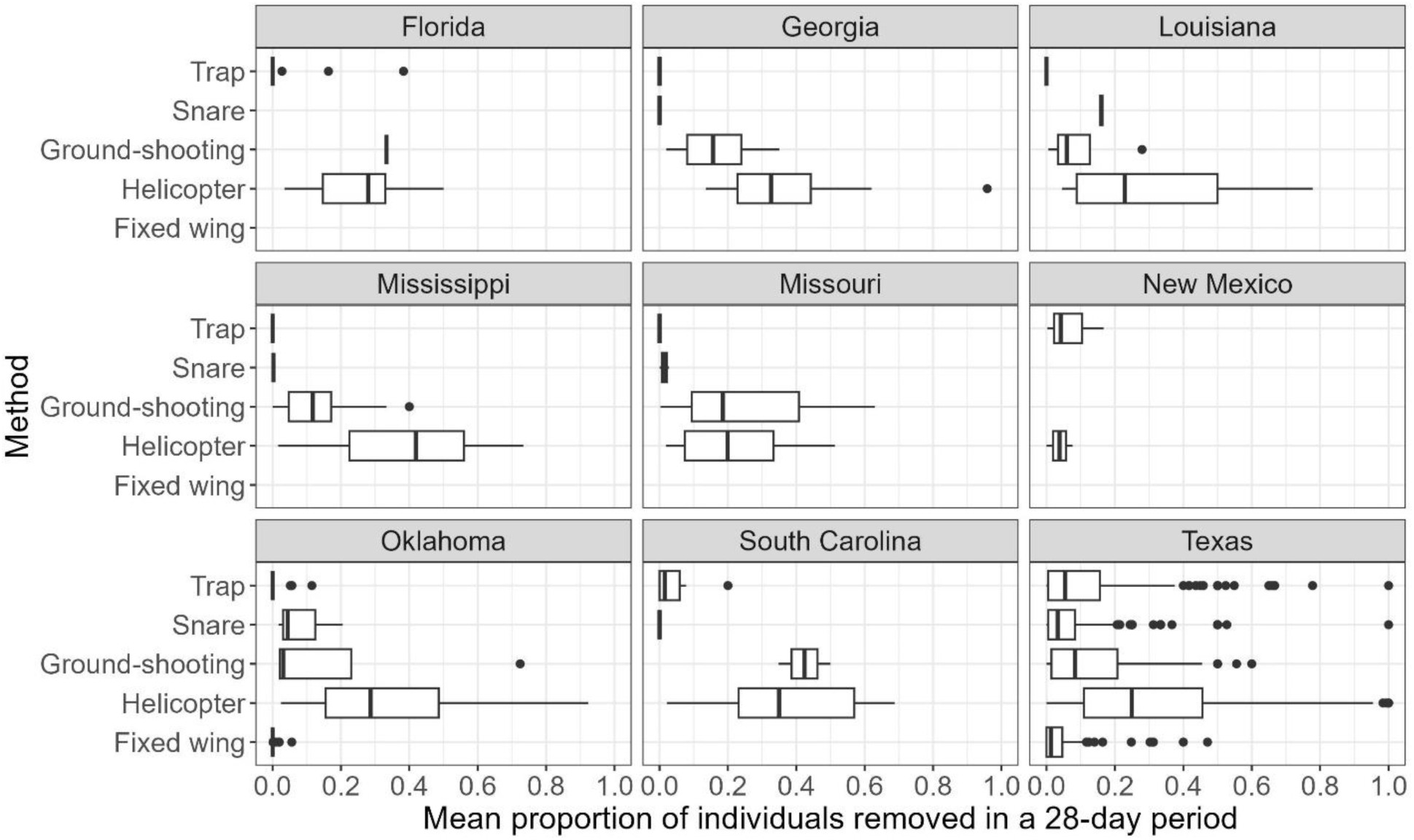
Summary of the average proportion of the population removed by each method at each property. Boxplots represent the median and interquartile range (IQR), where the whiskers extend plus or minus 1.5 times the IQR.

### Wild pig population growth rates vary in space

Accurately determining the population growth rates of invasive species is essential for effective management. This knowledge enables managers to forecast the speed of a species’ spread and the potential for ecosystem damage, allowing the implementation of targeted strategies that maximize population reduction and prevent ecosystem harm.

Individual populations exhibited a range of average growth rates. Across all properties the median of the average population growth rate ( 𝜆̅_𝑖_) was 0.96 (0 - 1.25), meaning over half (58%, n=570) of the properties had declining populations with ongoing control activities. There was also variation within and across states (Figure 2). For example, the median growth rate ranged from 0.5 to 1.03 for New Mexico and South Carolina, respectively. Additionally, there were nine states where at least half of their properties had a declining population (Appendix 1 Table S5).

**Figure 2.**
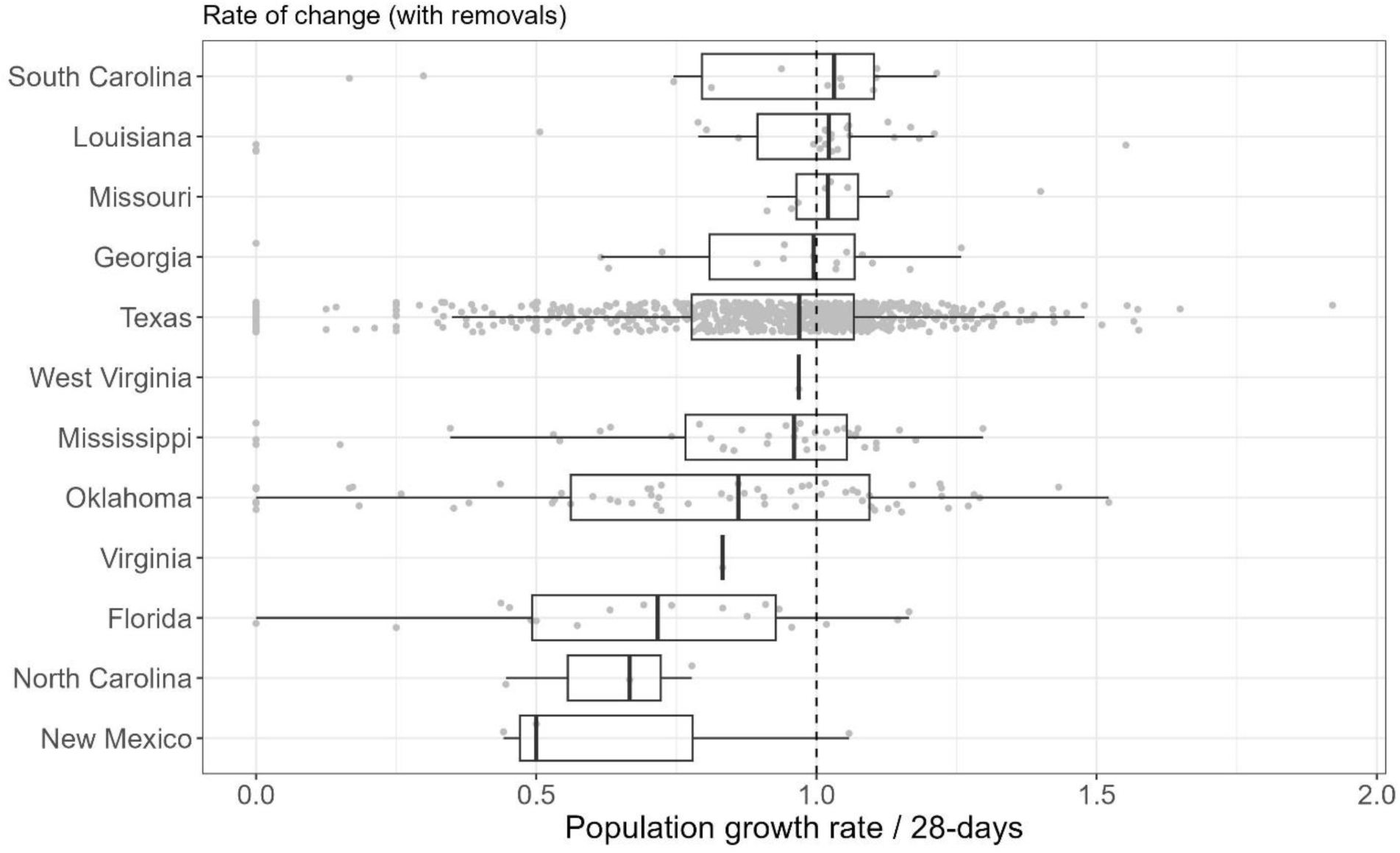
Each point is the average population growth rate for a single property. A growth rate greater than one indicates that a property’s population is, on average, growing. Values of zero represent properties that went extinct between the first and second observed primary periods. Boxplots represent the median and interquartile range (IQR), where the whiskers extend +/- 1.5 times the IQR.

Finally, there were 64 properties with a growth rate of zero, which were properties that eradicated wild pigs between their first and second primary periods with removal events. **Figure 2**

### Density is prevented due to management

Since the removal model inherently involves removing individuals for abundance estimation (Foster et al. 2025), simulating population dynamics with zero removals enables an assessment of management’s impact, and determines whether removals are ecologically and/or economically effective (Davis et al. 2018).

The variation in the effect of management was more pronounced as the initial density increased and as more time elapsed (Figure 3). For example, the amount of density prevented (i.e. the difference between the simulations without management compared to what happened with management) when the initial density of wild pigs was low (i.e. <1 pigs/km^2^) did not vary over five years, where after one year there was an estimated increase in density prevented by a median of 0.4 (4.8- −1.2) wild pigs/km^2^, and after five years there was no change at the median but the 95% C.I. showed anywhere from an increase of 3.4 to a decrease of 19 wild pigs/km^2^ prevented. However, when management occurred, the most likely outcome after five years in the low-density scenario was to keep populations stable. This contrasts with the no management simulations which were more likely to grow. Specifically, 75% of the low-density populations that experienced management grew by no more than 0.99 wild pigs/km^2^ while 25% of the simulations without management grew by at least 11.9 wild pigs/km^2^ (Figure 3).

**Figure 3:**
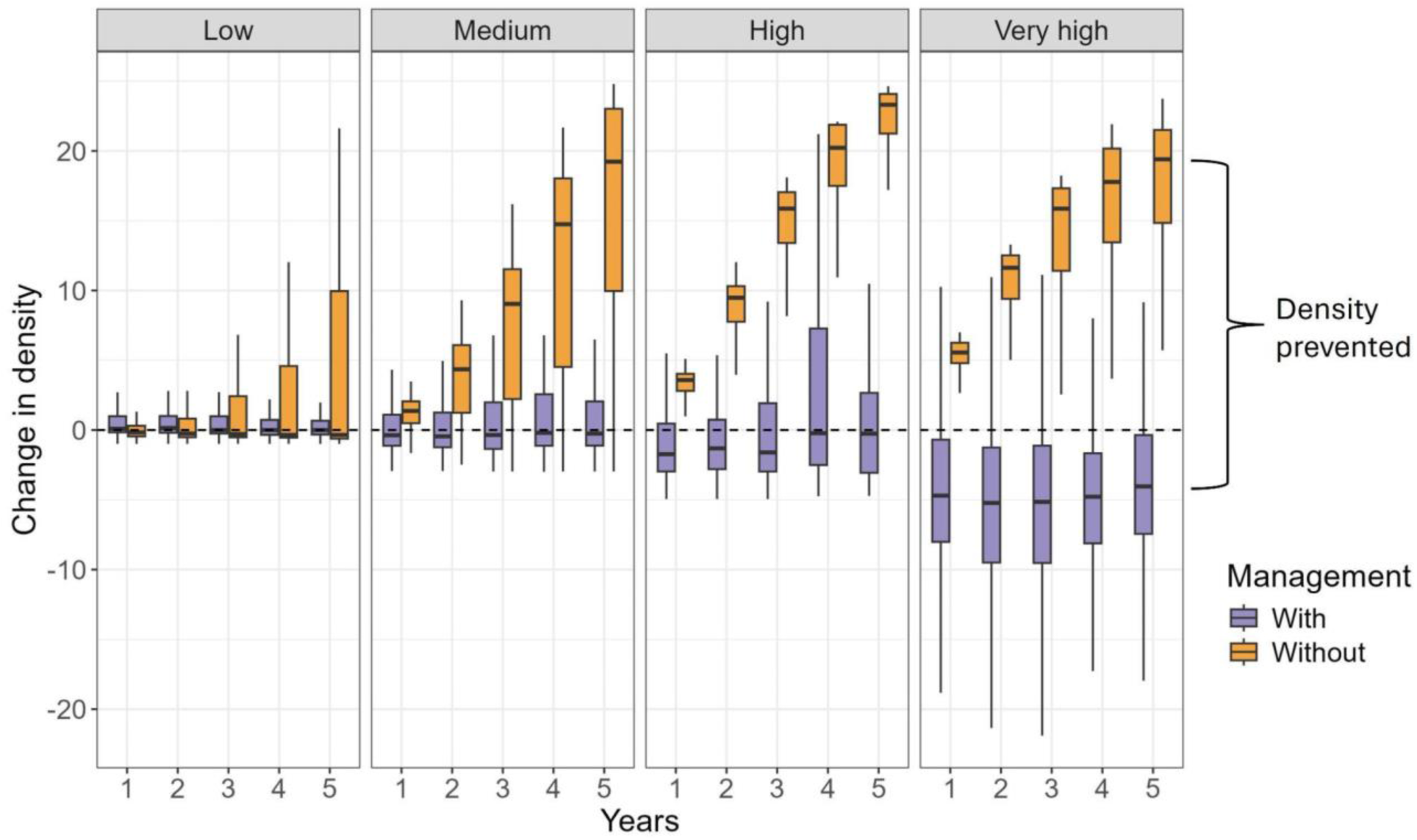
The change in median wild pig density relative to the initial median density at year 0 over a five-year period. The boxplots represent the median and interquartile range (IQR) of these changes, where the whiskers extend plus or minus 1.5 times the IQR with outliers removed. The Without management scenario (orange) is the result of the bootstrapping procedure, and the With management scenario (purple) compares median posterior density estimates at the specified interval to the initial posterior density estimate at year zero. The facets represent the initial density category, where Low is less than 1 wild pig/km^2^, Medium is between 1-3 wild pigs/km^2^, High is between 3-5 wild pigs/km^2^, and Very high is greater than 5 wild pigs/km^2^.

The amount of density prevented after one year increased with initial density, where 1.4 (−5.2-4.4), 4.7 (−8.2-8), and 9.6 (−6.2-21.5) wild pigs/km^2^ were prevented when initial density was medium (1-3 wild pigs/km^2^), high (3-5 wild pigs/km^2^), and very high (>5 wild pigs/km^2^), respectively. After five years, the amount of density prevented was similar across the medium, high, and very high initial density scenarios, where 17.7 (−3.5-25.4), 20.6 (−10.8-26.8), and 23.9 (−4.1-29.4) wild pigs/km^2^ were prevented, respectively. The absolute drop in density with management was most apparent in the very high-density properties, where the median reduction in density across all years was at least 4.24 wild pigs/km^2^ (Figure 3).

Additionally, across all properties, wild pig densities fell after five years at 44.1, 61, 56.9, and 80.3 percent of properties the low, medium, high, and very-high initial density scenarios, respectively. This contrasts with the simulations that did not conduct removals, where after five years wild pig densities fell at 60.8, 21.3, 0.0, and 0.0 percent of properties for the low, medium, high, and very high initial density scenarios, respectively.

### Population trajectories are modulated by effort and property size

The expected trajectory of the population (ETP) is a metric that combines the actual proportion removed (APR; Figure 1) with the population growth rate to estimate the net effect of removals on population size. This metric allows for direct comparison of removal effectiveness across properties and methods. To ease interpretation, we have categorized expected trajectories into three categories; slash (an expected reduction over 50%), shrink (an expected reduction between 0-50%), and grow (a population is expected to increase).

At the population level, on average, the most common EPT for each method except fixed wing aircraft was an expected shrinking of the population (Appendix 1 Tables S2-S4). The most common EPT for fixed wing aircraft was either growing or shrinking, which was expected for 53 populations for both EPTs. For every method except helicopters, slashing a population was, on average, the least likely EPT. For traps, snares, and ground-shooting, the most likely outcome was shrinking, then growing, and least likely was slashing.

Across all methods, the APR needed to slash a population was, on average, greater than to shrink a population by a factor of 2.72, and the APRs required to shrink a population were greater than those that resulted in a growing population by a factor of 3.7. For property area, across all methods, populations expected to grow were on properties 7.59 km^2^ larger than populations on properties expected to shrink, and populations expected to shrink were on properties 19.55 km^2^ larger than populations expected to slash.

Generalizing effort across methods is difficult because the unit of effort differs across methods (Table 1), and because each method is used in different ecological and management contexts (see Discussion). To attempt to generalize across methods (and therefore deployment context), we use a standardized metric of average total effort per unit deployed per area deployed (i.e. property area) to understand how the density of effort affects EPT. Here, the same pattern holds for all methods, where when the density of effort is greatest, the population was expected to slash, followed by shrink, and the lowest effort density leads to expected population growth (Figure 4C).

**Figure 4.**
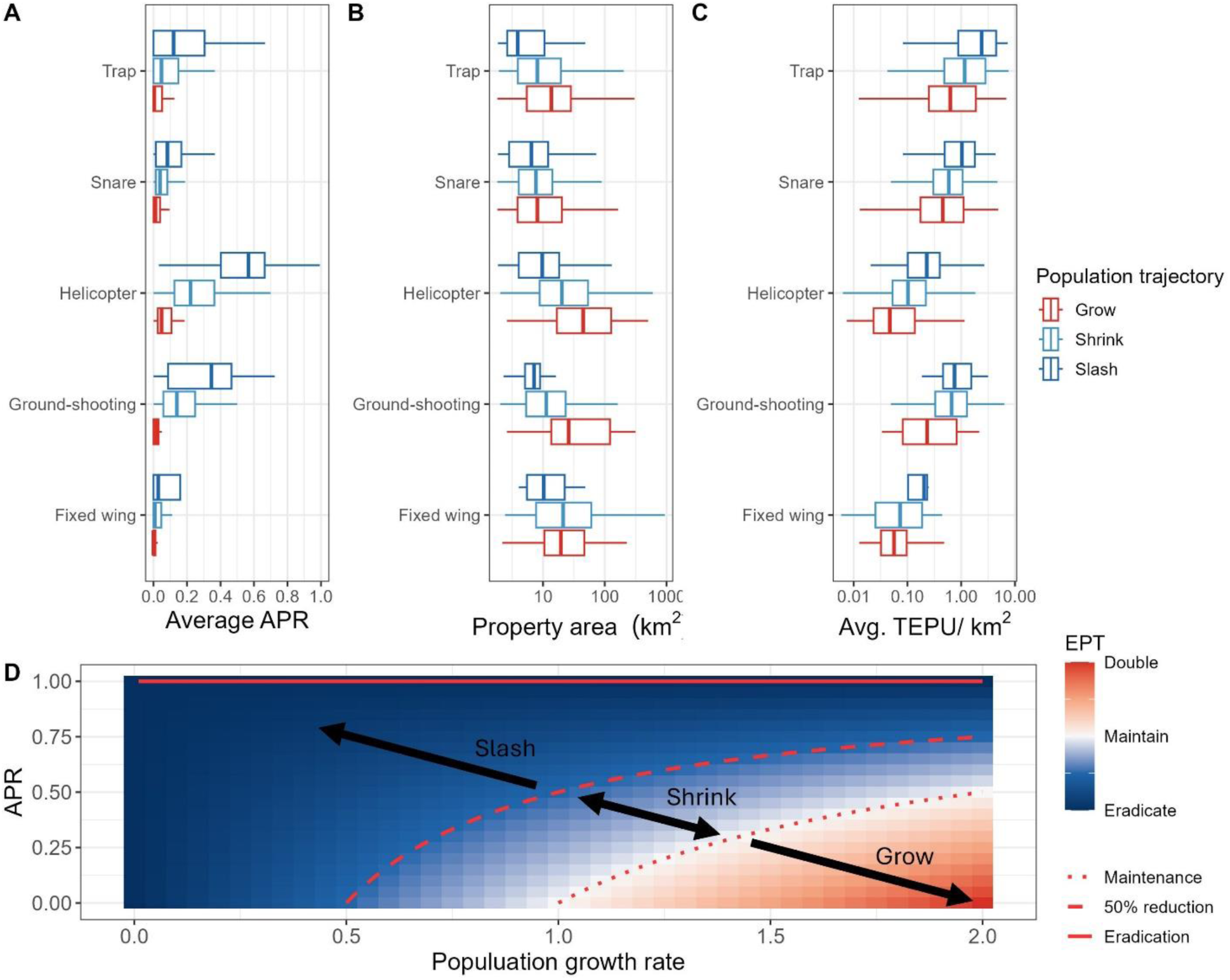
A) The average actual proportion of the population removed (APR) by each method and property. B) The area (km^2^) of each property where methods were used. C) The average total effort per unit (TEPU) deployed per unit area for each method in each property. D) The theoretically expected population trajectory (EPT) given growth rate and APR. The dashed line shows the threshold at which populations are expected to grow or increase, and the dotted line shows the threshold at which populations are expected to reduce by more (slash) or less (shrink) than half. For A-C, the data are grouped into the EPT given the average growth rate of the property where the method was deployed. Events that had an EPT of population growth are in red, events that had an EPT of a shrinking population are in light blue, and events that had an EPT of a slashing population are in dark blue.

**Table 1.** Summaries (median and 90% central interval) of the property-level average of the actual proportion of the population removed (APR), the area (km^2^) of the property where methods were deployed, and the property-level average of total effort per unit deployed for each method. Effort unit is the effort per unit deployed for each method. n is the number of properties that deployed a specific method.

| Method | APR | Area | Effort | Effort unit | n |
| --- | --- | --- | --- | --- | --- |
| Fixed wing | 0 (0-0.19) | 18.2 (3.4-157) | 1.3 (0.5-4.3) | Flight time (hours) per aircraft | 116 |
| Helicopter | 0.28 (0.03-0.74) | 17.2 (2.7-202.3) | 2.1 (0.7-7.9) | Flight time (hours) per helicopter | 429 |
| Ground-shooting | 0.1 (0-0.5) | 12.1 (2.4-143.1) | 8 (2-26.6) | Shooters per hour | 119 |
| Snare | 0.03 (0-0.24) | 7.5 (2.2-56.4) | 4.7 (1.6-9) | Snares per night | 464 |
| Trap | 0.03 (0-0.33) | 8.2 (2.2-92.8) | 10.3 (3.2-15.8) | Traps per night | 628 |

## Discussion

National-scale evaluation of invasive species management is challenging and rare (Ashander et al. 2022). While small studies are more prevalent, they are unlikely to achieve biodiversity goals at broad scales (McMillan et al. 2023). Here, we used a removal model that accurately predicts wild pig density to evaluate the effectiveness of five management methods at a national scale across 11 years of management. Results suggest that the effectiveness of wild pig management varies regionally and by removal method but is effectively reducing populations in most areas.

### Proportion of the population removed varies by method

When evaluating the effectiveness of various management techniques, it is crucial to consider the interplay between their logistical feasibility, cost implications, and overall impact on the population (Yokomizo et al. 2009). Traps represent the most frequently utilized method by USDA-APHIS-WS, being deployed on 63% of properties (Table 1). We found that traps yield the lowest returns in terms of the proportion of the population removed in a primary period, and that they are used on smaller properties than those of aerial operations (Table 1, Figure 1). Trap effectiveness on a catch-per-unit basis is affected by trap type (e.g. box, corral, etc.) and trigger type (e.g. remote or cellular operated) (Gaskamp et al. 2021; Lavelle et al. 2025), which is information we did not have and likely introduced bias in our estimates of individual capture rates by traps. However, given the large sample size of trapping events, we estimated likely population reductions on average for single bout of trapping regardless of trap type, trigger type, bait, or habitat (Foster et al. 2025). Additionally, traps remain a critical tool for wild pig management because of their low front end investment, ease of acquiring the necessary resources to begin trapping, ubiquitous applicability in different environmental conditions, and because they can be highly effective when they are applied persistently over a contiguous area for a long duration (i.e. 2 months; Snow et al. 2024).

Snares were the second-most utilized method, on 47% of properties, likely because they are usually the least expensive to deploy (Campbell and Long 2009). We found that snares had similar removal rates as traps (Table 1). Our previous work showed that snares (and traps) have diminishing returns on the area searched when more units are deployed (Foster et al. 2025), though the number of snares used may increase chances of capture and requires further investigation. The results for snares and traps highlight the importance of integrating other removal techniques as conditions change to maximize reduction potential.

Aerial methods (helicopter and fixed wing shooting) were used on a majority of properties, and helicopter-based gunning had the highest proportion of the population removed, which is in-line with previous work (Hamnett et al. 2024; Davis et al. 2018). This effectiveness is likely why helicopter-based gunning is the most cost-efficient (i.e. lowest cost per individual removed) method of wild pig management (Snow et al. 2024). However, the implementation of helicopter operations can be severely constrained by environmental factors, particularly dense forest cover and low population density, which limits when and where helicopters can be used (Smith et al. 2024; McCann et al. 2004). Logistical constraints on aerial units (helicopters and fixed-wing aircraft) include operational costs, limited flight time, regular maintenance, and pilots that have mandatory rest periods, which limit the continuous deployment of these resources. For very large properties (>221 km^2^, Appendix 1 Table S3), the EPT from aerial gunning is a growing population, likely because the cost and effort required for elimination increases as population density decreases (Choquenot et al. 1999), highlighting their limitation as a primary control method. Nevertheless, aerial gunning remains a primary tool for substantial population reductions, and the use of drones to aid helicopters in detecting individuals has been shown to reduce costs (Cross and Carlisle 2025; Cox et al. 2023).

Lastly, ground-shooting is the second-most effective method in terms of the proportion of the population removed but used on only 12% of properties (Table 1). Our estimate is most likely biased high because there is some amount of under reporting of unsuccessful ground-shooting. For example, USDA-APHIS-WS personnel might encounter a lone wild pig while conducting other activities and opportunistically remove the induvial and report it. Conversely, encountering this individual and not removing it will likely go unreported. Nonetheless, ground-shooting is often the least expensive removal method capable of achieving targeted results, including elimination (McCann and Garcelon 2008; Parkes et al. 2010; Snow et al. 2024; Hamnett et al. 2024). Additionally, ground-shooting is often highly effective in specific conditions, such as low population density, targeting lone individuals, or on properties that require greater control over operational aspects of removal such as urban or small properties.

While the methods evaluated above vary in their effectiveness individually, most removal programs operate using an integrated approach. Managers adjust methods as conditions and density change, such as helicopters for high-density populations, ground-shooting for targeted removals, and traps and snares because they are easier to implement. i.e. USDA-APHIS-WS tries to use the method that makes the most sense at the time, and most properties (60%) use more than one method to remove wild pigs.

### Wild pig population growth rates vary in space

Population growth rates significantly influence invasive species spread and establishment across a range of ecological systems (Miller et al. 2023). Here, we calculated the average population growth rate for wild pigs on 995 properties across the southeastern U.S. and showed that there was a large variation within and across states (Figure 2). While we focus on average growth rates, population growth rates are not static or uniform; they are inherently variable and influenced by a multitude of factors. For wild pigs this includes environmental conditions (Vetter et al. 2015), resource availability (Osada et al. 2015; Salinas et al. 2015), and human intervention such as translocation, agency-led removals, and hunting pressure (Hernández et al. 2018; Pepin, et al. 2017; Servanty et al. 2011).

Wild pig population dynamics are also influenced by unexplained variation attributed to environmental and/or demographic stochasticity and can be reactive to available forage conditions (Tabak et al. 2018; Miller 2017). Wild pigs evolved as pulsed resource consumers of mast; having larger litter sizes and reproducing more often under favorable forage conditions (Ostfeld and Keesing 2000). This variability in fecundity allows wild pig populations to rapidly grow under good forage conditions making them especially susceptible to transient population dynamics (Bieber and Ruf 2005; Miller 2017; Tabak et al. 2018). Nevertheless, the presence and magnitude of these processes can determine if a removal is compensatory or additive. If, for example, removals occur during a period of high natural mortality due to stochasticity, those removals would be additive (Kokko 2001; Sæther et al. 1996). Conversely, compensatory responses can limit the effectiveness of control efforts. For example, moderate and heavy harvest levels have produced similar population growth rates in wild pigs even though survival was lower in heavily harvested areas as increased immigration and reproduction likely offset those losses (Hanson et al. 2009).

### Density is prevented due to management

Evaluating the success of management efforts was significantly influenced by the initial density of the target population, where the magnitude of prevented density was largest in high-density areas (i.e. > 3 wild pigs/km^2^ Figure 3). This could be because population growth rates increase with initial population size in wild pigs (Miller et al. 2023; Tabak et al. 2018). Alternatively, when population density is high, management efforts tend to be more effective because individuals are easier to locate facilitating a more efficient application of control measures (Fischer et al. 2020; Tanadini and Schmidt 2011). However, our results indicate diminishing returns when initial density exceeds about 3 wild pigs/km², as density changes plateau in those scenarios (Figure 3). Although management actions are more effective when populations are at higher densities, increasing effort beyond that point yields progressively smaller benefits. The optimal level of management is the one that minimizes the combined costs of management and maximizes the impact management (Yokomizo et al. 2009).

However, we must also consider the density-related objectives of management strategies. Properties with high densities typically have the objective of mitigating damage rather than elimination (Gaskamp et al. 2018). This approach is more appropriate for pervasive or widespread populations where complete elimination is impractical, and the goal shifts to reducing their impact to an acceptable level (Green and Grosholz 2021). Furthermore, controlling populations when they are at their peak density is the most cost-effective strategy even when elimination is unlikely, as might be the case in a dense, localized population (Davis et al. 2018; Keiter et al. 2017; Baxter et al. 2008).

In contrast to high density areas, we found little effect of management when the initial density was low (Figure 3). Furthermore, evaluating the true impact of management at low densities is difficult because its effect may be too weak or obscured by other ecological factors such as immigration or stochasticity (Hanson et al. 2009). Additional unintended effects could be due to the management activities themselves such as in increase in density because bait was used during a removal campaign and thus increased food availability (Ditchkoff et al. 2017). Furthermore, management activities could have been conducted on populations already in decline due to unstable sex ratios (i.e. male dominated) and thus have fluctuating growth rates (Koons et al. 2006). This suggests that what appears to be a slight positive effect of management at low densities might, in fact, be a natural phase of declining population growth rate, irrespective of intervention (Cerini et al. 2023). These factors make the inference on population growth rates at low densities difficult, and thus the attribution of the observed changes in density solely to management actions unreliable, and highlights the importance of robust baselines such as density estimates pre removal (Snow et al. 2025), which we did not have in this study.

Nevertheless, a key question that remains is whether management has any effect when populations are at low densities. If the population shows no measurable response at low density despite intervention, this may indicate the strategy is ineffective or that other factors dominate. In such cases, resources might be better directed toward higher-density areas. On the other hand, low-density areas might need more resources to accomplish elimination, which suggests that either the current level of resources is insufficient for this goal or that removal activities cease once agricultural damage has fallen to an acceptable level. However, properties with low densities, which are often areas where invasive species have recently established, will likely have the objective of elimination as the probability of achieving elimination is highest at the beginning of an invasion (Simberloff 2003; Venette et al. 2021). Early intervention can also reduce elimination effort requirements (Buhle et al. 2012), and reducing low-density populations to levels that do not allow for population growth increases elimination probability (Liebhold et al. 2016). However, due to the high reproductive potential of wild pigs, elimination attempts that fail to remove all individuals can result in reestablishment (Snow et al. 2020).

Finally, the apparent negligible difference between managed and unmanaged areas might be because the environmental conditions or natural regulatory processes were already sufficient to keep the population regulated, rendering management efforts redundant or minimally impactful. This suggests that removals at low densities tend to be compensatory. Furthermore, we observed that properties that had more continuous management efforts can contribute to better outcomes, which raises questions about resource allocation and cost-effectiveness. An important consideration is the long-term consequence of inaction. If a population is left uncontrolled, there is not only an accumulation of individuals but also a compounding of associated damages or ecological impacts (Bobek et al. 2017).

### Population trajectories are modulated by effort and property size

What happens to a population after removals varies by a populations’ growth rate, which can be used to determine the threshold proportion of the population one must remove to stop a population from growing (Hone et al. 2010; Hone 2001). Here, we showed how to predict the management outcome on a relative scale given population growth and individuals removed. For example, for a desired outcome (i.e. EPT), equation 4 could be used to determine how many individuals to remove and which removal method to use to achieve the desired goal. We have shown that effectiveness in wild pig management is highly dependent on factors like property size, effort, and the method used for removals (Figure 4).

Effective management of animal populations, particularly invasive species, presents significant logistical challenges such as the availability of resources (which varies by removal method) that often limit the achievable level of population reduction (Taylor and Hastings 2004; Hastings et al. 2006; Pepin et al. 2020). Additionally, as the managed area becomes larger, achieving a substantial population reduction becomes increasingly difficult. This suggests that the effort required to manage larger areas does not scale linearly from what is required of smaller areas (Pepin et al. 2017), as the largest population reductions occur when effort per unit area is maximized (Bengsen et al. 2022). And, while removal efforts are often targeted to meet specific needs, there is an upper boundary on what is feasible and how much equipment, resources, and time is available. On small properties this upper bound is likely never reached leading to more effective management. But as property size increases this upper bound is likely reached and there is only so much reduction that is logistically feasible. This may explain why the success of ungulate removal campaigns is more likely on small properties (Snow et al. 2025).

There appears to be an optimal allocation of units and time for searching a given property size (Figure 4), and an increase in effort has been shown to increase the detection of invasive species at low abundances (Mehta et al. 2007), and increase spatial optimization (Nishimoto et al. 2021). However, in some cases we show that removals can still lead to an expected increase in abundance because removals do not offset gains from reproduction and/or immigration (Figure 4). Indeed, wild pigs can recover from a 68% reduction in population size in as little as three months (Garabedian and Kilgo 2024), but they can also fall in response to sustained management over a 3-year period (Treichler et al. 2023).

The conventional definition of effort in wildlife population models (removal models being just one example) often oversimplifies the complexities involved, requiring a more nuanced understanding that incorporates available resources and logistical constraints (Koch et al. 2020). The concept of effort could explicitly include the practical limitations imposed by human resources, equipment availability, and operational logistics. For example, simply stating “trap nights” as an estimator for effort can be misleading without considering the underlying model of personnel and other constraints that enable those trap nights. In fact, Snow et al. (2024), reported that trapping was the most physically demanding removal method compared to aerial operations and toxic baiting. The logistical constraint of how many aircraft can be flown at once, or how many traps can be deployed and managed simultaneously, highlight the need to account for real-world operational ceilings. Quantifying logistic constraints (e.g. personnel hours, fuel consumption, equipment deployment, budgets, and maintenance downtime) in a formalized way may help increase the accuracy of the estimate of the area searched, and thus the accuracy of the prediction of the proportion of the population removed. Additionally, the influence of logistical constraints on removal can help identify those aspects that need additional research or resources to improve removal efficiency. In turn, this could increase the chance of achieving desired management outcomes on large properties. This would be advantageous, and may offset the cost of removing the last individuals of a population which are by far the most expensive to remove (Fischer et al. 2020).

While population models often fail to incorporate the constraints on effort, they do frequently include environmental factors (St. Clair et al. 2013; Rivera and McCrea 2021). We suggest that there is an underlying function that determines the actual effort applied. This could be in the form of a truncated distribution to model the upper bound of effort given logistical constraints, as truncated distributions improve parameter estimates in Bayesian hierarchical models by enabling more accurate inference from incomplete data and better uncertainty quantification (Cangelosi and Hooten 2009). The concept of observer effects in mark-resight and distance models serves as a good corollary for the unquantified limitations on effort (Conn and Alisauskas 2018; Link and Sauer 1997). Just as observer bias can affect detection probabilities, logistical constraints can similarly impact the effectiveness of management and monitoring interventions (Witmer 2005).

Better spatial information, such as mapping exact removal locations, could significantly improve understanding of how each method searches a property. Although we acknowledge that doing so at the scale of this study is impractical. Furthermore, a significant data gap exists regarding the distribution of traps and snares, such as whether they were opportunistically, randomly, or systematically placed, which can bias population estimates (Whysong and Miller 1987). This impacts the interpretation of effort and success rates, especially for snares where a large number are deployed. Further research is needed to identify constraints with the greatest negative effects on management effort.

## Conclusion

Here, we developed a method to evaluate management progress for a national program targeted at one of North America’s most invasive species. We incorporated the five most prevalent methods of removal to estimate the proportion of the population each method removes in a single bout of effort and found that helicopters are the most effective and traps are used most often. Additionally, wild pig density declined across most properties managed by USDA-APHIS-WS over the last 11 years, especially when initial population density was high. Our results suggest that with sustained management activities, populations should decline in the long term on USDA-APHIS-WS managed properties. We also showed that effort does not scale with property size, and that additional effort is required to effectively manage large properties. Recognizing these limitations is essential for creating realistic models and ensuring sustainable invasive species management, as it helps allocate resources effectively to maximize success (Giljohann et al. 2011).

## Acknowledgments

We thank Michael Glow and Tim Smyser for feedback on the manuscript. This research was supported in part by an appointment to the Animal and Plant Health Inspection Service (APHIS) Research Participation Program administered by the Oak Ridge Institute for Science and Education (ORISE) through an interagency agreement between the U.S. Department of Energy (DOE) and the U.S. Department of Agriculture (USDA). ORISE is managed by ORAU under DOE contract number DE-SC0014664. All opinions expressed in this paper are the author’s and do not necessarily reflect the policies and views of USDA, DOE, or ORAU/ORISE.

