## Appendix 1 for "The effect of long-term management on wild pig (*Sus scrofa × domesticus*) populations across the southeastern United States"

### 1. Methods

#### 1.1 Density prevented without management

The model described in (Foster et al. 2025) does not include density dependence, because the regulating factor on the populations were the management actions themselves (i.e. the number of individuals removed $C_{it}$, eq. S1). The discrete population growth rate ($\lambda_i$) can be reduced to the additive effects of survival ($\phi$) and per capita reproduction ($\frac{\zeta}{2}$) (eq. S2). Meaning that, without management actions the population model is an exponential growth model (eq. S3). As such, we modified the population model to include density dependence to stop runaway populations due to the exponential growth where carrying capacity at each property ($K_i$) was set to 30 individuals/km$^2$ (Pepin et al. 2017), and we reparametrized growth in terms of the intrinsic growth rate ($r_i = \lambda_i - 1$, (Skalski et al. 2005)) (eq. S4).

$$N_{it} = (N_{it-1} - C_{it})\phi + (N_{t-1} - C_{it})\ \frac{\zeta}{2} \quad \text{[S1]}$$

$$N_t = (N_{it-1} - C_{it})(\phi + \frac{\zeta}{2}) \quad \text{[S2]}$$

$$N_{it} = N_{it-1}\lambda_i \quad \text{[S3]}$$

$$N_{it} = N_{it-1} + N_{it-1}r_i(1 - \frac{N_{it-1}}{K_i}) \quad \text{[S4]}$$

#### 1.2 Expected population trajectory

The formulation for the expected population trajectory (EPT) is as follows. Starting with the general exponential growth equation to predict abundance $N$ at time $t$,

$$N_t = (N_{t-1} - C_t)\lambda \quad \text{[S5]}$$

where $\lambda$ is the instantaneous growth rate and is equal to the sum of survival and per capita reproduction and $C$ is the number of individuals removed from the population (eqs. S1 and S2). Our goal is to determine the expected trajectory of the population represented as a proportion of the current population size. Therefore, $N_{it} = (1 + EPT)N_{it-1}$ where $EPT$ is the expected population trajectory. In this formulation if the population is expected to double then $EPT$ = 1, if no change is expected then $EPT$ = 0, and if eradication is expected then $EPT$ = -1. We can substitute in our new value for $N_{it}$ on the lefthand side of equation S5 to get

$$(1 + EPT)N_{t-1} = (N_{t-1} - C_t)\lambda \,. \quad [S6]$$

Next, we can express the number of individuals removed as a proportion of the current population size ($C_t = APR * N_{t-1}$; where $APR$ is bound between zero and one) to get

$$(1 + EPT)N_{t-1} = (N_{t-1} - (APR * N_{t-1}))\lambda \,. \quad [S7]$$

Then, solving for $EPT$;

$$(1 + EPT)N_{t-1} = \lambda N_{t-1}(1 - APR) \quad [S8]$$

$$1 + EPT = \lambda(1 - APR) \quad [S9]$$

$$EPT = \lambda(1 - APR) - 1 \quad [S10]$$

**1.3 Iterative Bayesian model fitting**

There was a total of 11 rounds of iterative updates, where the initial 105 properties described in Foster et al. (2025) represent the first round. Each of the 890 remaining properties were randomly assigned to one of the subsequent rounds. For each round the MCMC was run across seven chains until burn-in occurred (i.e. a PSRF value < 1.1) and each parameter reached a minimum of 1000 effective samples (Gelman and Rubin 1992). The number of properties added by round, the total number of properties fit in each round, and the total number of MCMC iterations per round are reported in Appendix 1 Table S1.

For rounds 2-11, the posterior distribution from the previous round was used as the prior for the subsequent one. Specifically, we used moment matching to approximate posterior parameter distributions into hyperparameters of the priors. To approximate the Gaussian posterior distributions (on the log scale) for the priors of $\rho_k$ (how effort scales with area searched by method $K$), $\omega_k$ (the degree of overlap in searched area for each trap and snare deployed), $\gamma_k$ (saturating constant of traps and snares), and $\nu$ (per capita reproduction), we calculated the mean and precision. $\beta_k$ (the effect of landscape covariates on detection probability for each method $K$) was also approximated with a Gaussian distribution. For the prior on $\phi_\mu$(mean survival rate per primary period), we calculated the mean ($\mu_\phi$) and variance ($\tau_\phi$) of the previous rounds posterior, then calculated shape $\alpha_\phi = \mu(\frac{\mu(1-\mu)}{\tau} - 1)$ and rate $\beta_\phi = (1-\mu)(\frac{\mu(1-\mu)}{\tau} - 1)$ which became the hyperparameters for the Beta distribution on $\phi_\mu$. For the prior on $\psi$ (shrinkage rate for survival), we calculated the mean ($\mu_\psi$) and variance ($\tau_\psi$) of the previous rounds posterior, then calculated the shape $\alpha_\psi = \frac{\mu^2{}_\psi}{\tau_\psi}$ and rate $\beta_\psi = \frac{\mu_\psi}{\tau_\psi}$ which became the hyperparameters for the Gamma distribution on $\psi$. How the posterior distributions from the initial fit from (Foster et al. 2025) compare to the posteriors after the 11$^{th}$ round are in Appendix 1 Figures S1-S4 and Table S6.

### 2. Results

**Table S1**

| Round | n properties added | n properties total | MCMC iterations | Burn-in |
|---|---|---|---|---|
| 1 | 105 | 105 | 65,000 | 18,237 |
| 2 | 109 | 214 | 115,000 | 18,443 |
| 3 | 89 | 303 | 160,000 | 96,021 |
| 4 | 100 | 403 | 170,000 | 105,420 |
| 5 | 81 | 484 | 140,000 | 44,835 |
| 6 | 94 | 578 | 150,000 | 45,036 |
| 7 | 95 | 673 | 150,000 | 18,045 |
| 8 | 70 | 743 | 230,000 | 23,046 |
| 9 | 74 | 817 | 110,000 | 72,618 |
| 10 | 84 | 901 | 230,000 | 18,447 |
| 11 | 94 | 995 | 115,000 | 11,546 |

**Table S1** - Summary of the number of properties added during the iterative fitting procedure. Round is the iterative fitting iteration, n properties added is the number of properties added in each round, n properties total is the total number of properties fit each round, MCMC iterations is the number of total iterations each round ran and Burn-in is the burn-in iteration.

**Table S2 - Proportion of population removed and expected population trajectory**

| Method | Expected population trajectory | Proportion of the population removed |
|---|---|---|
| Fixed wing | Grow | 0 (0-0.05) |
| Fixed wing | Shrink | 0.01 (0-0.22) |
| Fixed wing | Slash | 0.03 (0-0.41) |
| Ground-shooting | Grow | 0.01 (0-0.1) |
| Ground-shooting | Shrink | 0.14 (0-0.41) |
| Ground-shooting | Slash | 0.35 (0-0.65) |
| Helicopter | Grow | 0.04 (0-0.23) |
| Helicopter | Shrink | 0.22 (0.03-0.5) |
| Helicopter | Slash | 0.54 (0.14-0.89) |
| Snare | Grow | 0.01 (0-0.11) |
| Snare | Shrink | 0.04 (0-0.17) |
| Snare | Slash | 0.08 (0-0.26) |
| Trap | Grow | 0 (0-0.17) |
| Trap | Shrink | 0.05 (0-0.27) |
| Trap | Slash | 0.12 (0-0.65) |

**Table S2 -** Summary (median and central 90% CI) by method of the proportion of the population removed associated with each expected population category. Grow means the population is expected to grow after removals take place, shrink means the population is expected to fall by at most 50%, and slash indicates a population is expected to fall by more than 50% after removals.

**Table S3 - Property area and expected population trajectory**

| Method | Expected population trajectory | Property area ($km^2$) |
|---|---|---|
| Fixed wing | Grow | 19.38 (4.05-131.12) |
| Fixed wing | Shrink | 18.45 (3.39-284.25) |
| Fixed wing | Slash | 11.73 (4.35-43.89) |
| Ground-shooting | Grow | 43.79 (3.19-163.28) |
| Ground-shooting | Shrink | 12.55 (2.54-77.78) |
| Ground-shooting | Slash | 7.11 (2.37-18.45) |
| Helicopter | Grow | 44.47 (4.89-345.48) |
| Helicopter | Shrink | 20.88 (3.53-233.42) |
| Helicopter | Slash | 9.89 (2.36-51.6) |
| Snare | Grow | 8.5 (2.17-93.89) |
| Snare | Shrink | 7.69 (2.26-49.27) |
| Snare | Slash | 6.07 (2.21-20.23) |
| Trap | Grow | 14.16 (2.37-141.64) |
| Trap | Shrink | 8.25 (2.27-68.24) |
| Trap | Slash | 4.05 (1.97-26.71) |

**Table S3** - Summary (median and central 90% CI) by method of the property areas ($km^2$) associated with each expected population category. Grow means the population is expected to grow after removals take place, shrink means the population is expected to fall by at most 50%, and slash indicates a population is expected to fall by more than 50% after removals.

**Table S4 - Effort and expected population trajectory**

| Method | Expected population trajectory | TEPU/km$^2$ |
|---|---|---|
| Fixed wing | Grow | 0.05 (0.02-0.39) |
| Fixed wing | Shrink | 0.07 (0.01-0.29) |
| Fixed wing | Slash | 0.21 (0.04-0.24) |
| Ground-shooting | Grow | 0.23 (0.03-1.61) |
| Ground-shooting | Shrink | 0.64 (0.08-3.34) |
| Ground-shooting | Slash | 0.91 (0.28-4.02) |
| Helicopter | Grow | 0.05 (0.01-0.25) |
| Helicopter | Shrink | 0.1 (0.01-0.49) |
| Helicopter | Slash | 0.2 (0.03-0.68) |
| Snare | Grow | 0.44 (0.04-2.7) |
| Snare | Shrink | 0.59 (0.06-2.06) |
| Snare | Slash | 1.03 (0.11-3.93) |
| Trap | Grow | 0.6 (0.06-5.11) |
| Trap | Shrink | 1.13 (0.11-4.73) |
| Trap | Slash | 2.36 (0.23-6.54) |

**Table S4** - Summary (median and central 90% CI) by method of the effort per unit deployed associated with each expected population category. The average total effort per unit deployed (TEPU/km$^2$) is the effort per unit deployed is the effort (hobbs hours per aircraft for fixed wing and helicopters, shooters per hour for ground-shooting, and the number of traps or snares deployed per night for traps and snares) per unit area. Grow means the population is expected to grow after removals take place, shrink means the population is expected to fall by at most 50%, and slash indicates a population is expected to fall by more than 50% after removals.

**Table S5**

| State | Mean | SD | n |
|---|---|---|---|
| Missouri | 1.06 | 0.15 | 8 |
| West Virginia | 0.97 | NA | 1 |
| Louisiana | 0.91 | 0.38 | 26 |
| Georgia | 0.90 | 0.31 | 15 |
| South Carolina | 0.88 | 0.33 | 12 |
| Texas | 0.87 | 0.32 | 802 |
| Virginia | 0.83 | NA | 1 |
| Mississippi | 0.83 | 0.33 | 39 |
| Oklahoma | 0.79 | 0.39 | 61 |
| Florida | 0.70 | 0.31 | 18 |
| New Mexico | 0.67 | 0.34 | 3 |
| North Carolina | 0.63 | 0.17 | 3 |

**Table S5** - The mean and standard deviation (SD) of average property growth rates across properties in each state (n is the number of properties in each state).

**Figure S1**

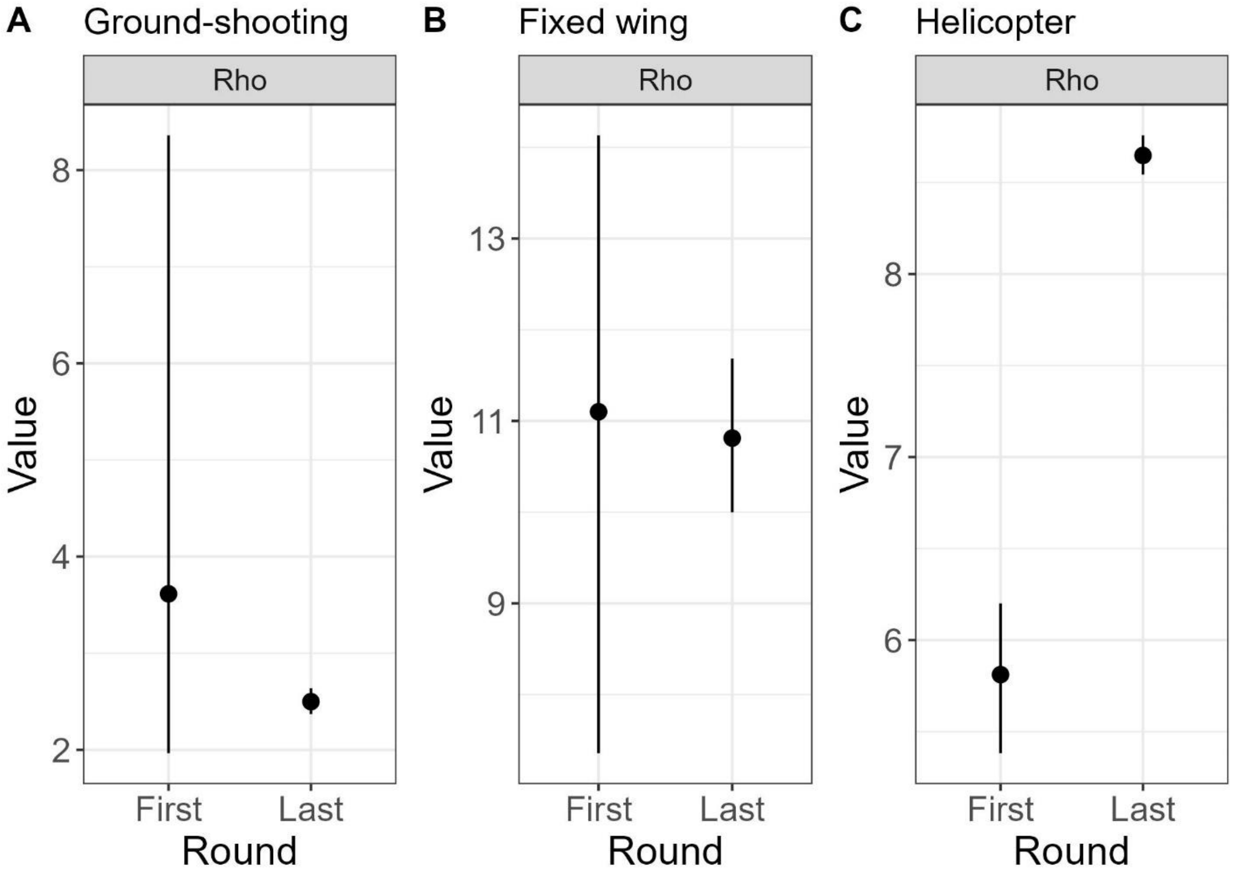


**Figure S1** – Posterior distributions (mean and 95% CI) after the first and last round of fitting the model for the scaling coefficient ($\rho$) on effort (to determine area searched) for the active removal methods (equation 4 Foster et al. (2025)).

**Figure S2**

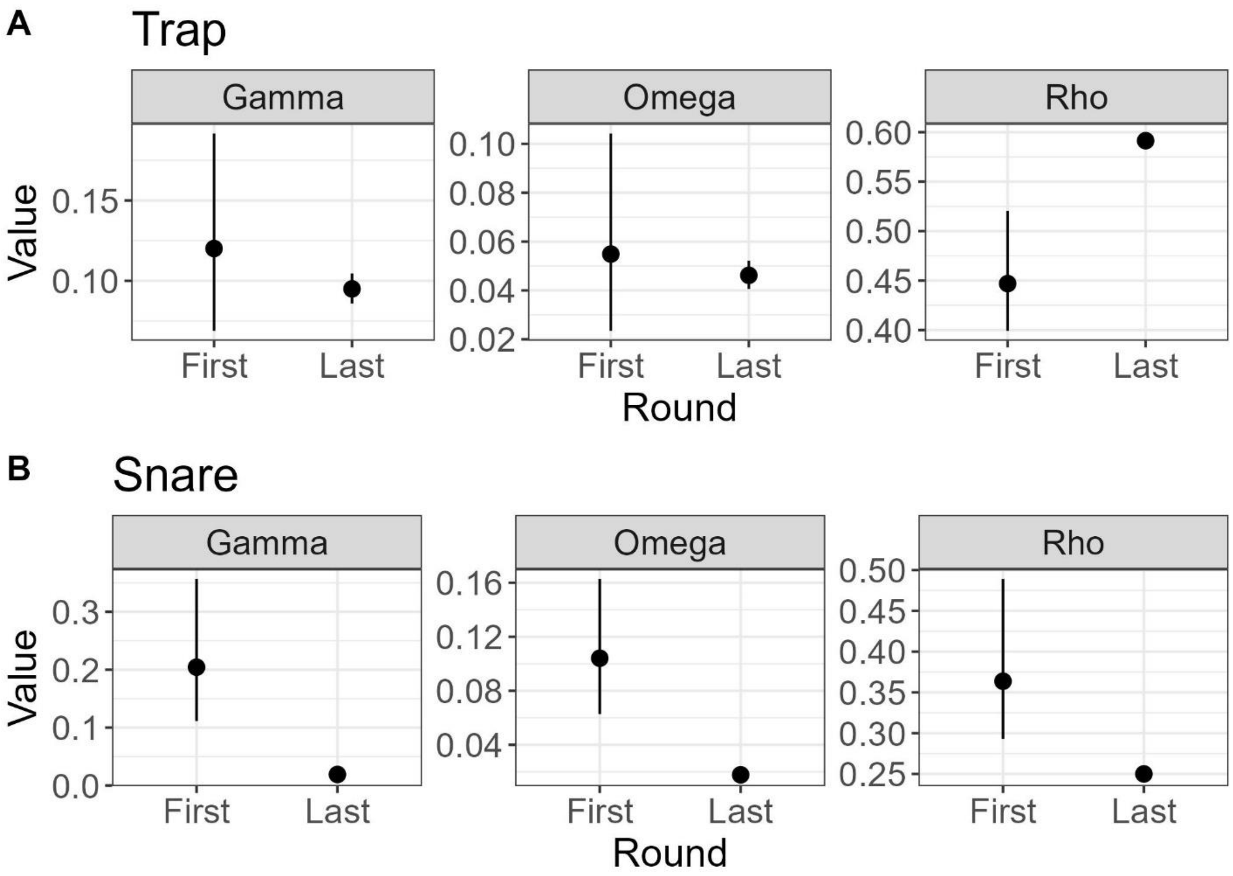


**Figure S2** – Posterior distributions (mean and 95% CI) after the first and last round of fitting the model for the saturating constant ($\gamma$), amount of overlap ($\omega$), and scaling coefficient ($\rho$) on effort (to determine area searched) for the passive removal methods (equation 5 Foster et al. (2025)).

**Figure S3**

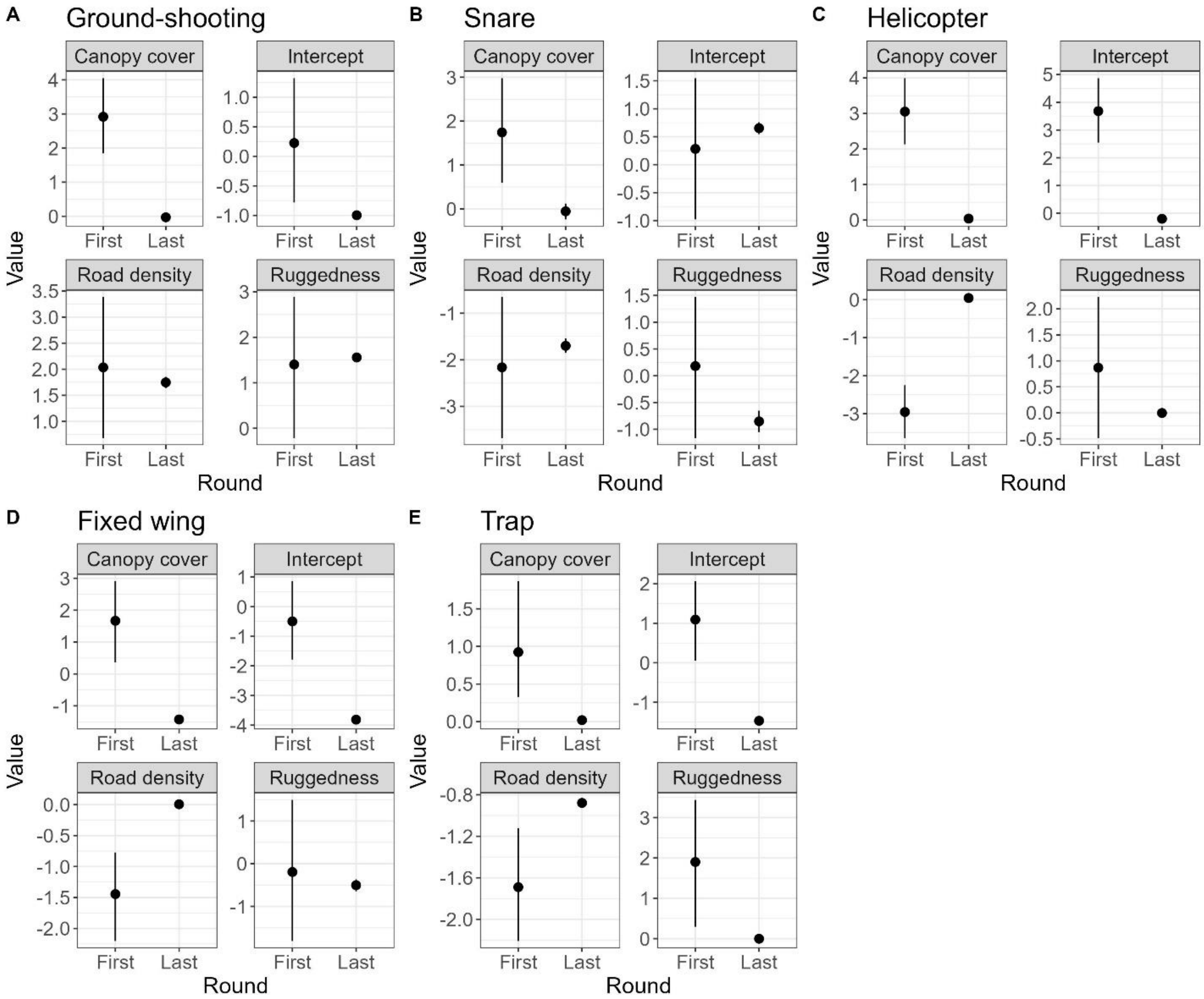


**Figure S3 -** Posterior distributions (mean and 95% CI) after the first and last round of fitting the model for the method specific coefficients on land cover (to determine removal probability) for the all removal methods (equation 6 Foster et al. (2025)).

**Figure S4**

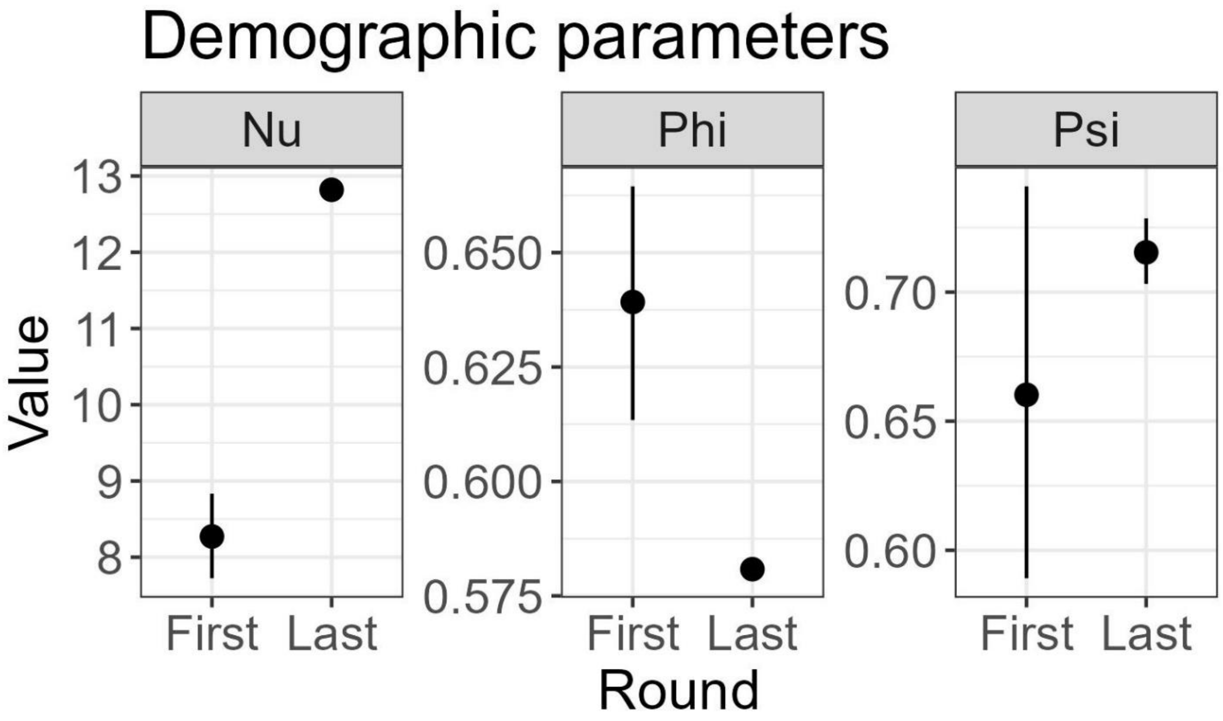


**Figure S4 -** Posterior distributions (mean and 95% CI) after the first and last round of fitting the model for the per capita reproduction ($\nu$), 28-day survival rate ($\phi$), and shrinkage on survival ($\psi$) (equation 1 Foster et al. (2025)).

**Figure S5**

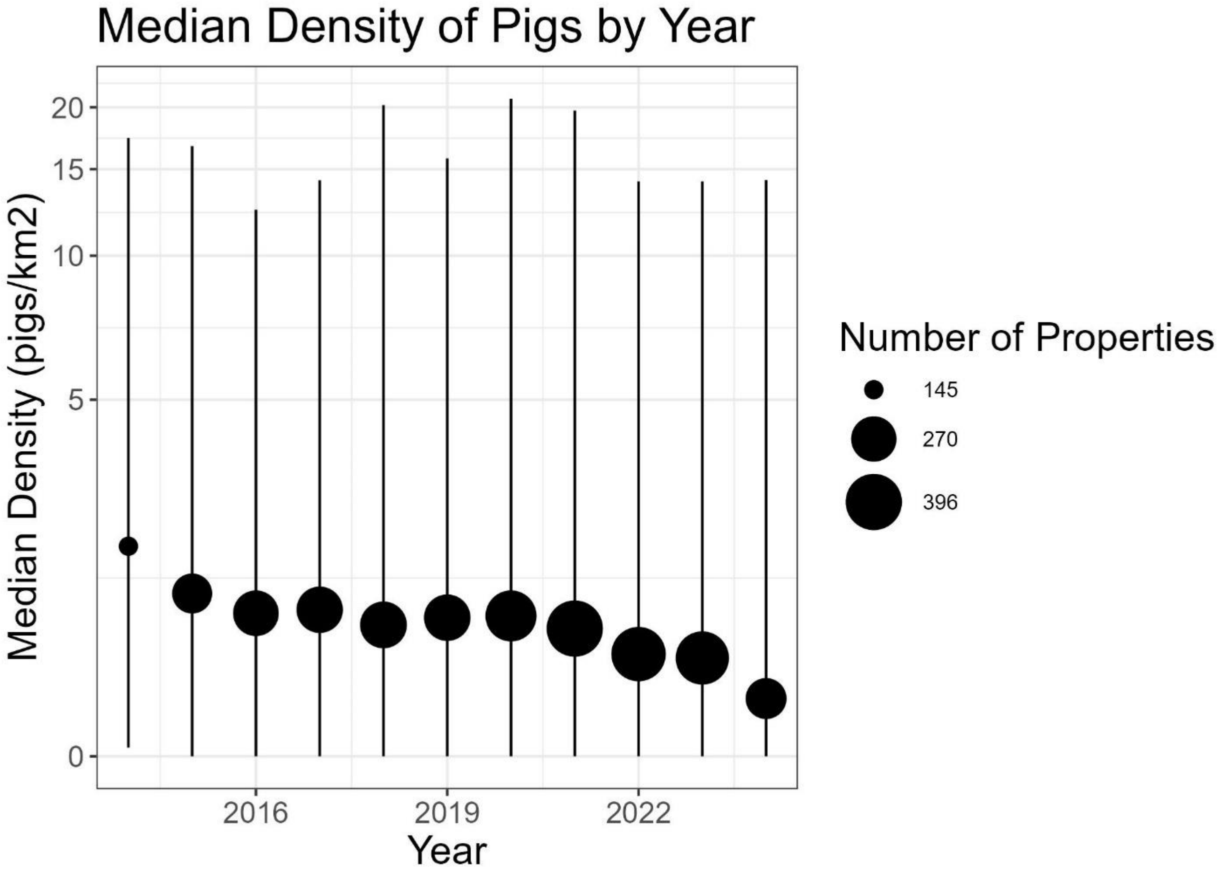


**Figure S5 –** The distribution (median and central 90% interval) of median posterior density estimates across the 12 states and 11 years where USDA-WS removals occurred (n=12,432 individual primary periods from 2014-01-28 to 2024-06-30 across 995 properties). Point size represents the number of properties with density estimates each year.

**Table S6**

| Parameter | Method | Land | First Round | Last Round |
|---|---|---|---|---|
| $\beta_1$ | Ground-shooting | Intercept | 0.25 (0.54) | -1 (0.035) |
| $\beta_2$ | Fixed wing | Intercept | -0.49 (0.68) | -3.81 (0.046) |
| $\beta_3$ | Helicopter | Intercept | 3.69 (0.59) | -0.2 (0.011) |
| $\beta_4$ | Snare | Intercept | 0.29 (0.64) | 0.65 (0.054) |
| $\beta_5$ | Trap | Intercept | 1.09 (0.51) | -1.47 (0.02) |
| $\beta_{1,1}$ | Ground-shooting | Road density | 2.03 (0.69) | 1.75 (0.053) |
| $\beta_{1,2}$ | Ground-shooting | Ruggedness | 1.39 (0.79) | 1.56 (0.053) |
| $\beta_{1,3}$ | Ground-shooting | Canopy cover | 2.92 (0.57) | -0.03 (0.032) |
| $\beta_{2,1}$ | Fixed wing | Road density | -1.45 (0.37) | 0.01 (0.036) |
| $\beta_{2,2}$ | Fixed wing | Ruggedness | -0.18 (0.84) | -0.5 (0.073) |
| $\beta_{2,3}$ | Fixed wing | Canopy cover | 1.65 (0.65) | -1.43 (0.046) |
| $\beta_{3,1}$ | Helicopter | Road density | -2.95 (0.36) | 0.04 (0.014) |
| $\beta_{3,2}$ | Helicopter | Ruggedness | 0.86 (0.7) | 0 (0.018) |
| $\beta_{3,3}$ | Helicopter | Canopy cover | 3.05 (0.47) | 0.04 (0.011) |
| $\beta_{4,1}$ | Snare | Road density | -2.16 (0.77) | -1.7 (0.079) |
| $\beta_{4,2}$ | Snare | Ruggedness | 0.17 (0.68) | -0.85 (0.102) |
| $\beta_{4,3}$ | Snare | Canopy cover | 1.75 (0.61) | -0.06 (0.092) |
| $\beta_{5,1}$ | Trap | Road density | -1.68 (0.27) | -0.88 (0.015) |
| $\beta_{5,2}$ | Trap | Ruggedness | 1.89 (0.79) | 0 (0.029) |
| $\beta_{5,3}$ | Trap | Canopy cover | 0.97 (0.39) | 0.02 (0.015) |
| $\gamma_1$ | Snare | NA | 0.22 (0.08) | 0.02 (0.002) |
| $\gamma_2$ | Trap | NA | 0.12 (0.04) | 0.1 (0.006) |
| $\nu$ | NA | NA | 8.27 (0.34) | 12.82 (0.027) |
| $\omega_1$ | Snare | NA | 0.11 (0.03) | 0.02 (0.001) |
| $\omega_1$ | Trap | NA | 0.06 (0.02) | 0.05 (0.004) |
| $\phi_\mu$ | NA | NA | 0.64 (0.02) | 0.58 (0.001) |
| $\psi_\phi$ | NA | NA | 0.66 (0.05) | 0.72 (0.008) |
| $\rho_1$ | Ground-shooting | NA | 4.31 (1.88) | 2.5 (0.067) |
| $\rho_2$ | Fixed wing | NA | 10.99 (1.8) | 10.82 (0.434) |
| $\rho_3$ | Helicopter | NA | 5.81 (0.21) | 8.65 (0.054) |
| $\rho_4$ | Snare | NA | 0.38 (0.08) | 0.25 (0.002) |
| $\rho_5$ | Trap | NA | 0.45 (0.04) | 0.59 (0.004) |

**Table S6** – Posterior distribution summaries (mean and standard deviation) for parameters in the model after the first round of fitting the model and the last round of fitting the model. $\beta$ parameters are intercept and method specific coefficients tied to land cover to determine capture probability. $\gamma$ is the saturating constant for traps and snares. $\nu$ is per capita reproduction. $\omega$ is the amount of overlap in search area for traps and snares. $\phi_{mu}$ and $\psi_{\phi}$ are mean 28-day survival rate and shrinkage for survival, respectively. $\rho$ is the scaling factor on effort for each method to determine search area.
